# Space and epigenetic inheritance determine inter-individual differences in siderophore gene expression in bacterial colonies

**DOI:** 10.1101/2023.08.21.554085

**Authors:** Subham Mridha, Tobias Wechsler, Rolf Kümmerli

## Abstract

Heterogeneity in gene expression among cells in clonal groups is common in bacteria. Albeit ubiquitous, it often remains unclear what the sources of variation are and whether variation has functional significance. Here, we tracked the expression of genes involved in the synthesis of iron-chelating siderophores (pyoverdine and pyochelin) in individual cells of the bacterium *Pseudomonas aeruginosa* during colony growth on surfaces using time-lapse fluorescence microscopy, to explore potential sources and functions of cellular heterogeneity. Regarding sources, we found that the physical position of cells within the colony and epigenetic gene expression inheritance from mother to daughter cells significantly contributed to cellular heterogeneity. In contrast, cell pole age and cellular lifespan had no effect. Regarding functions, our results indicate that cells optimize their siderophore investment strategies (pyoverdine vs. pyochelin) along a gradient from the centre to the edge of the colony. Moreover, we found evidence that cell lineages with above-average siderophore investment increase the fitness of cell lineages with below-average investment through cooperative sharing of secreted siderophores. Altogether, our study highlights that single-cell experiments combined with automated image and cell-tracking analyses can identify sources of heterogeneity and yield adaptive explanations for gene expression variation among clonal bacterial cells.

## Introduction

It is commonly observed that bacteria in a group show variation in their behaviour albeit being clonal and living in the same environment^1–7^. Both intrinsic and extrinsic factors can contribute to individual heterogeneity. At the extrinsic level, individual cells might experience differences in their micro-environment, which can spur differences in their molecular activities and thus trigger variation in their behavioral responses^8–10^. For example, cells growing at the edge of a biofilm face different environmental conditions than cells at the interior of the biofilm^11^. At the intrinsic level, heterogeneity can arise because biological processes such as gene expression and protein synthesis are noisy^1^. Random noise plays an important role, but there are also deterministic factors such as cell age^12,13^ or epigenetic inheritance^9,14^ that can contribute to cellular heterogeneity. Although heterogeneity is ubiquitous in microbial systems, it is often hard to identify its underlying sources, and it is even harder to assess whether the observed heterogeneity is beneficial for the bacteria and reflects an evolved adaptive strategy^5^.

Here, we explore both putative sources and adaptive functions of cellular heterogeneity in the expression of two siderophore synthesis genes in clonal colonies of the bacterium *Pseudomonas aeruginosa*. Siderophores are secondary metabolites secreted by bacteria to scavenge iron from the environment^15,16^. While genes involved in the synthesis of the two siderophores pyoverdine and pyochelin are heterogeneously expressed in *P. aeruginosa*, little is known on the underlying sources of heterogeneity^17–19^. Moreover, siderophore molecules are secreted and their function (iron-acquisition) is shared between cells in a group, leading to several possible adaptive explanations for heterogeneity^7^. We use fluorescent time-lapse microscopy to simultaneously track cellular heterogeneity in the expression of two genes encoding enzymes involved in the synthesis of pyoverdine and pyochelin. We start with single cells placed on agarose pads and track cell identity, spatial positioning, cell division events, and siderophore gene expression over five hours (every 10 minutes). Cells divide five to eight times during this time frame and form a single-layer colony. The tracking of cellular heterogeneity through space and time allows us to differentiate between extrinsic and intrinsic sources of variation, and to test adaptive hypotheses. We focus on four potential sources of heterogeneity and two adaptive explanations. Below, we formulate specific hypotheses for each source and adaptive explanation.

Extrinsic factor – the micro-environment. *P. aeruginosa* (like other bacteria) senses iron limitation and secretes siderophores to scavenge this essential trace element from the environment^20–22^. Previous work revealed sophisticated regulatory mechanisms that allow bacteria to sense both iron limitation and the rate of incoming iron-loaded siderophores to adjust siderophore synthesis in a fine-tuned manner^23–26^. Due to this high sensitivity, we hypothesize that cells will sense differences in iron and siderophore concentrations in their micro-environment depending on their spatial position in the colony (e.g., center versus edge) and adjust their siderophore gene expression accordingly.

Intrinsic factor A – cell life span. In growing colonies, individual cells will divide at different time points, leading to heterogeneity in a cell’s life span. We hypothesize that variation in life spans can translate into heterogeneity in siderophore gene expression. For example, cells with longer life spans might show higher gene expression than cells that divide more quickly. Intrinsic factor B – genealogy. Mother cells can pass on their gene expression status to their daughter cells^9^. We hypothesize that this form of epigenetic inheritance results in gene expression levels to be more similar between sister cells than between non-related cells in the group. This hypothesis thus assumes that gene expression noise is propagated through genealogical lineages. Intrinsic factor C – cell pole age. Upon binary cell division, each bacterium has an old and a new pole (Fig. S1). The old pole is inherited from the mother cell, while the new pole is formed upon division. During the next cell division, one of the daughter cells inherits the cell pole of the grandmother, while the other daughter cell inherits the pole of the mother. Like this, relative cell pole age can be tracked through time. We hypothesize that variation in cell pole age will translate into gene expression heterogeneity^12^. For instance, cellular functions might change with age such that cells with older cell poles show different siderophore gene expression levels than cells with younger cell poles.

Adaptive explanation A – cost-to-benefit optimization. Previous work revealed that the costs and benefits differ for pyoverdine and pyochelin^19,27,28^. While pyoverdine has a higher affinity for iron than pyochelin (K_a:_ 10^32^ M^−1^ vs. 10^18^ M^−2^)^29^, it is more costly to make. We previously showed that a plastic dual siderophore investment strategy is most beneficial for *P. aeruginosa*, whereby the bacteria predominantly invest in pyoverdine under stringent iron limitation and switch to increased pyochelin investment under moderate iron limitation^28^. Assuming that cells can sense variation in iron availability in their micro-environment, we hypothesize that cells optimize their pyoverdine vs. pyochelin gene expression strategy depending on their spatial position in the colony. Adaptive explanation B – public goods production to help clonemates. Pyoverdine and pyochelin are secreted in the environment. They can deliver iron to other cells than the producer and are therefore considered public goods, accessible to other colony members^30^. Given this shared function, we hypothesize that cells with low individual fitness (i.e., cells with slow division rates, old cells) can indirectly increase group fitness via a disproportionally high siderophore investment to improve iron nutrition and growth of clonemates.

## Results

### Colony-level siderophore gene expression varies in response to iron limitation and time

We exposed cells of *P. aeruginosa* PAO1 to either moderate iron limitation (casamino acids medium, CAA, characterized by naturally low iron content) or stringent iron limitation (CAA supplemented with 400μM of the synthetic iron chelator 2-2′-bipyridyl). We used a variant of *P. aeruginosa* PAO1 that featured a double gene expression reporter construct. Specifically, the reporter strain *PAO1pvdA::mcherry–pchEF::gfp* had the promotors of the *pvdA* gene (encoding a pyoverdine synthesis enzyme) and *pchEF* genes (encoding two pyochelin synthesis enzymes) fused to *mcherry* and *gfp* genes, respectively. The reporter construct is stably integrated as a single copy into the chromosome^19^.

In a first step, we conducted colony-level analysis to obtain an overview on the siderophore gene expression of *P. aeruginosa* in a spatially structured environment over time and in response to iron limitation (Fig. 1). As expected, we found that pyoverdine gene expression was significantly higher under stringent iron limitation than under moderate iron limitation (Fig 1a+b, F_1,43_ = 42.24, p < 0.0001). Pyoverdine gene expression significantly declined over time in both media (Fig 1c, moderate iron limitation: *t*_18_ = -5.82, p < 0.0001; stringent iron limitation: *t*_25_ = -10.77, p < 0.0001). Similar patterns were observed for pyochelin gene expression, which was significantly higher under stringent versus moderate iron limitation (Fig 1d+e, F_1,43_ = 25.59, p < 0.0001). Pyochelin gene expression also significantly declined with time (Fig 1f, moderate iron limitation: *t*_18_ = -3.82, p = 0.0025; stringent iron limitation: *t*_25_ = -2.31, p = 0.0296) although the drop is less pronounced than for pyoverdine gene expression.

**Fig 1.**
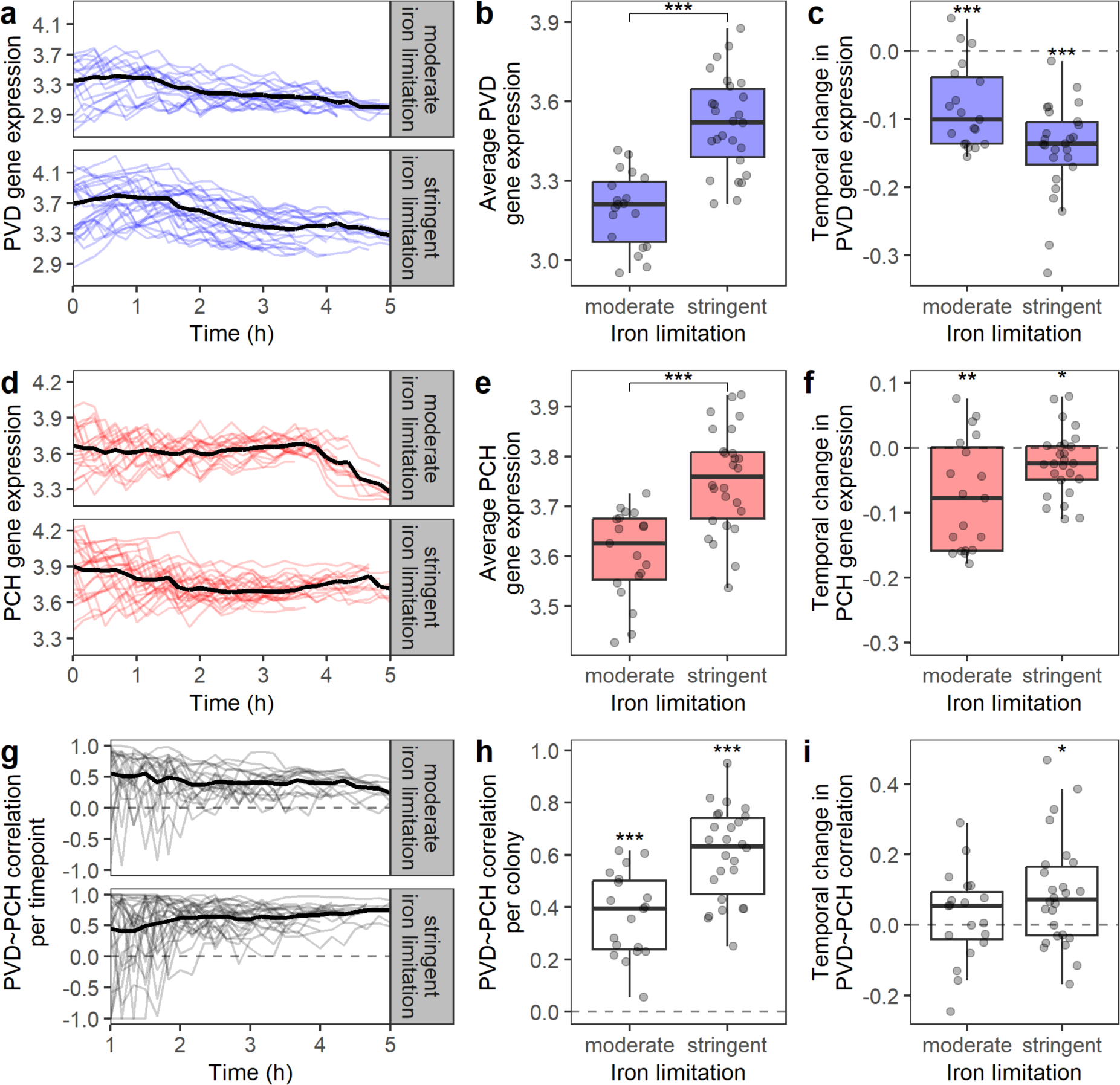
Patterns of siderophore gene expression in growing colonies of *P. aeruginosa* in response to different levels of iron limitation. (**a+d**) Pyoverdine (blue) and pyochelin (red) gene expression per colony (mean across all cells) under moderate (CAA + 0μM bipyridyl, N = 19) and high iron limitation (CAA + 400μM bipyridyl, N = 26). Black lines indicate the average values across all colonies. (**b+e**) Average pyoverdine and pyochelin gene expression per colony (shown as individual dots). (**c+f**) Temporal change in pyoverdine and pyochelin gene expression, measured as the slope of the linear regression between time and gene expression within a colony. (**g**) Temporal patterns of gene expression correlations between pyoverdine and pyochelin (mean grey value) among cells within a colony. (**h**) Average gene expression correlation between pyoverdine and pyochelin across all time points with colony size ≥ 16 cells. (**i**) Temporal change in the gene expression correlation between pyoverdine and pyochelin, measured as the slope of the linear regression between time and pyoverdine and pyochelin correlation coefficients per colony. Boxplots show the median and the interquartile range (IQR), while whiskers indicate minimum and maximum values. * p < 0.05, ** p < 0.01, *** p < 0.001.

We then asked whether pyoverdine and pyochelin gene expression are correlated at the individual cell level. For this analysis, we considered all time points during which colonies contained at least 16 cells because correlation coefficients varied stochastically between -1 and +1 in very small colonies (Fig. 1g). We found that pyoverdine and pyochelin gene expression were positively correlated across cells within a colony (Fig 1h, moderate iron limitation: *t*_18_ = 10.14, p < 0.0001; stringent iron limitation: *t*_25_ = 17.50, p < 0.0001), whereby the positive association was significantly stronger under stringent iron limitation compared to moderate iron limitation (Fig. 1h, ANOVA: F_1,43_ = 20.75, p < 0.0001). The strength of the positive association significantly increased over time under stringent iron limitation (*t_2_*_5_ = 2.92, p = 0.0145), but did not change under moderate iron limitation (*t*_18_ = 0.81, p = 0.4280) (Fig. 1i).

In summary, our colony-level analyses confirm results from liquid-culture experiments^19^, showing that (i) *P. aeruginosa* cells respond to variation in iron limitation by adjusting siderophore gene expression, and (ii) pyoverdine and pyochelin gene expression is positively correlated across cells, but less so under moderate iron limitation.

### Spatial gradients of siderophore gene expression activities within colonies

We hypothesized that gradients of iron availability exist within colonies, which will manifest in cells adjusting their siderophore gene expression depending on their spatial position within the colony. To test this hypothesis, we measured the Euclidean distance of each cell from the edge of the colony and correlated its siderophore gene expression with this distance (Fig. S2). We calculated the spatial gene expression correlation for each time point imaged and grouped the extracted values into four distinct classes of colony size (Fig 2a + b).

**Fig 2.**
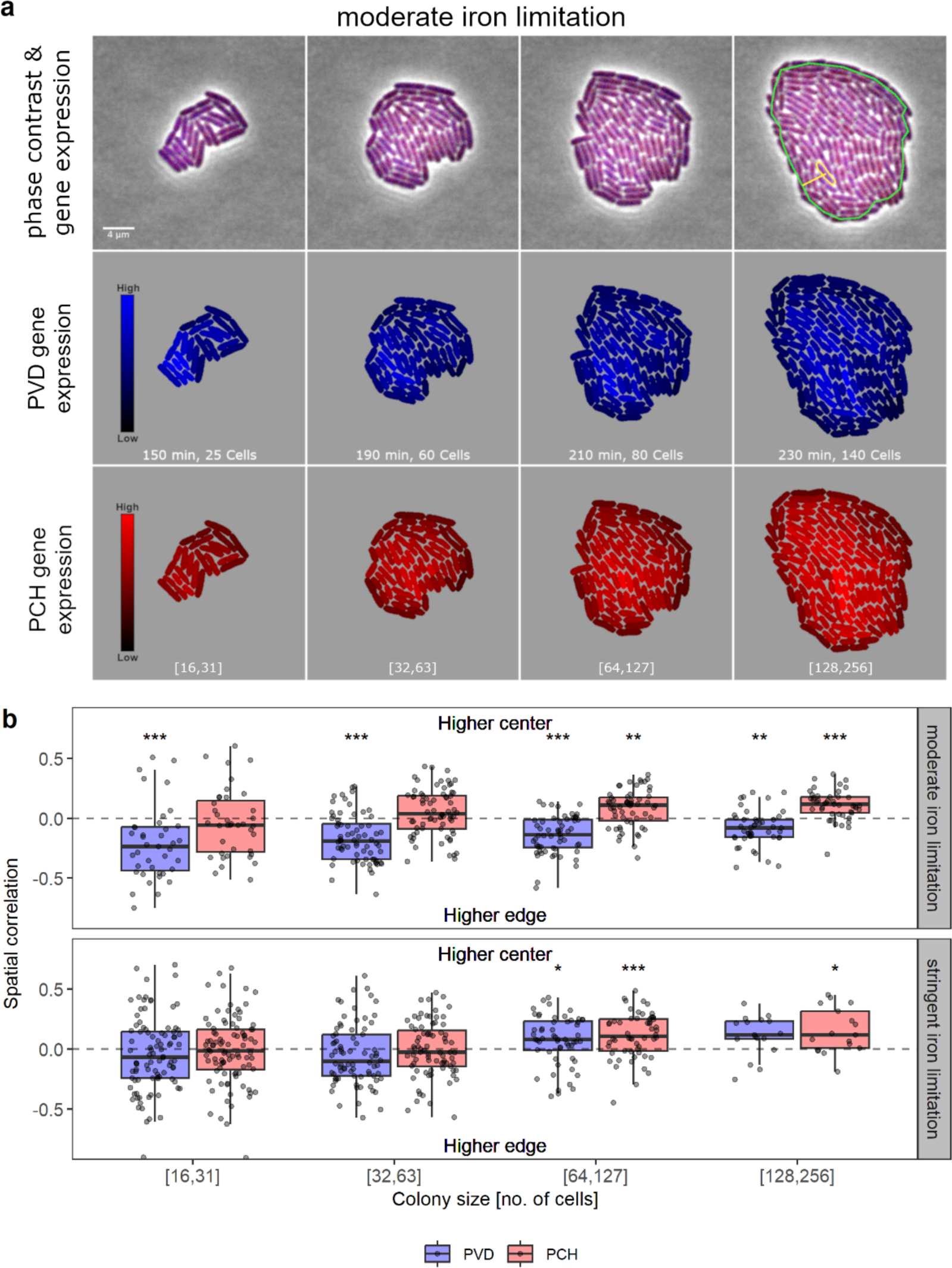
Spatial gradients in gene expression arise under moderate iron limitation. (**a**) Snapshots of a representative colony over time under moderate iron limitation (CAA + 0μM bipyridyl). Top row shows the overlay of phase contrast, GFP (pyochelin, blue) and mCherry (pyoverdine, red) channels. The green line indicates the edge of the colony and the yellow line the distance to the edge for an individual cell. Middle and bottom rows show segmented cells with heatmap colors indicating pyoverdine (blue) and pyochelin (red) gene expression levels, respectively. (**b**) Spatial correlation coefficients between pyoverdine (blue) or pyochelin (red) gene expression and the distance of cells from the colony edge calculated per colony and timepoint and grouped by colony size category. Boxplots show the median and the interquartile range (IQR), while whiskers indicate minimum and maximum values. Asterisks indicate significant differences from the dashed line. * p < 0.05, ** p < 0.01, *** p < 0.001.

Under moderate iron limitation, we found significant spatial gradients, whereby pyoverdine gene expression significantly increased in cells that were closer to the colony edge (Fig. 2b top panel, Table S1). The relationship held for all colony sizes, but the strength of the association declined in larger colonies (ANOVA: F_3,447_ = 6.26, p = 0.0004). We observed the opposite pattern for pyochelin, for which gene expression significantly declined towards the colony edge (Fig. 2b top panel, Table S2) with the association becoming stronger in larger colonies (ANOVA: F_3,447_ = 8.49, p < 0.0001). Spatial gradients were much weaker under stringent iron limitation. In this environment, cells closer to the center of the colony tend to invest more in both siderophores – pyoverdine (Fig 2b bottom panel) and pyochelin (Fig 2b bottom panel) – but the effects only became significant in larger colonies (Table S1+S2).

### Cell lifespan does not correlate with siderophore gene expression

We hypothesized that variation in the lifespan of cells (defined as time between two cell divisions) could be an intrinsic factor contributing to variation in siderophore gene expression. To test our hypothesis, we first created lineage trees for each colony, where nodes represent individual cells and branch length reflect their lifespans (Fig 3a shows a representative example). We then calculated the average siderophore gene expression per branch length and correlated this metric to the lifespan of the cell.

**Fig 3.**
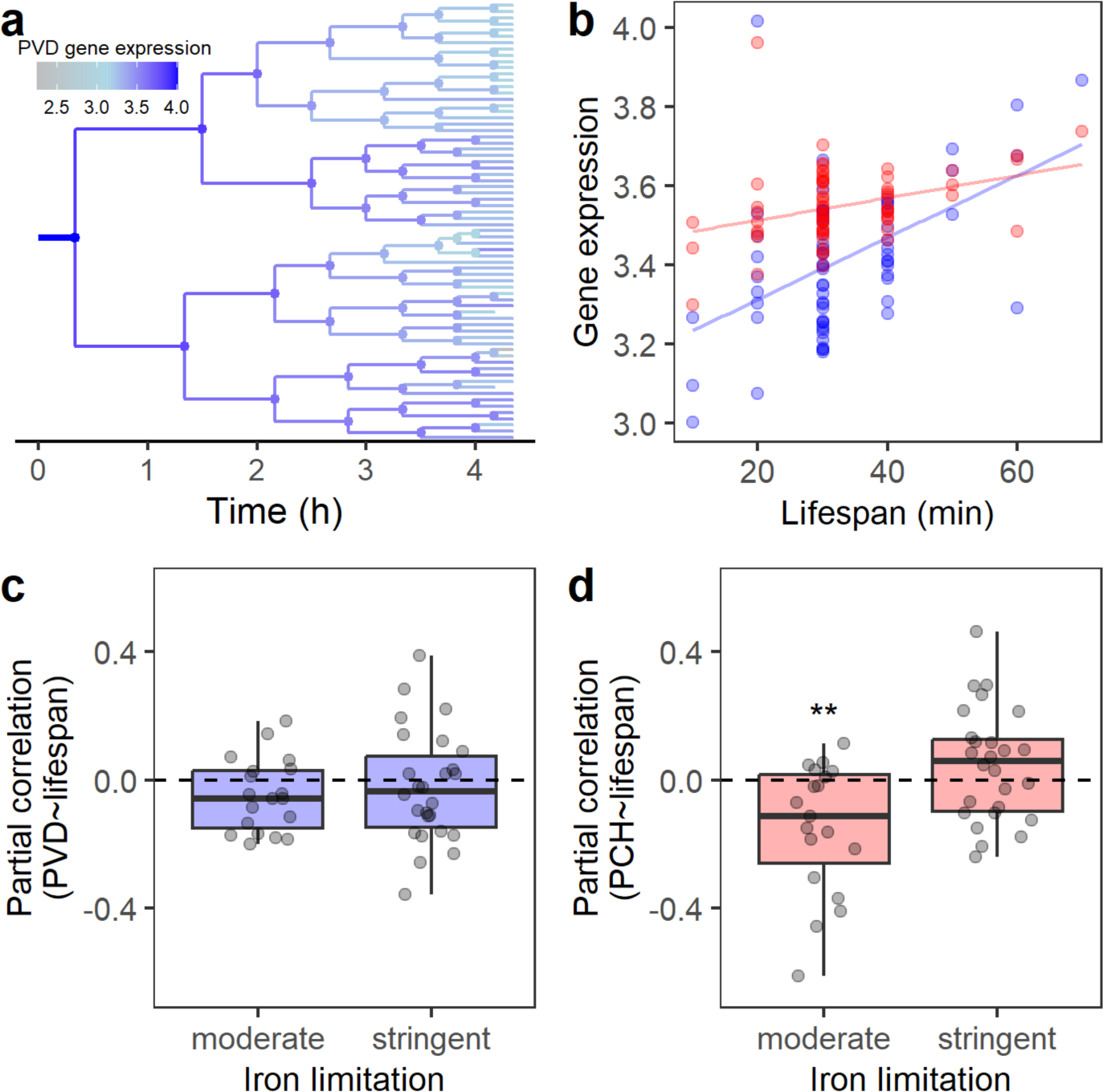
No positive associations between lifespans of cells and siderophore gene expression. (**a**) Lineage tree of a colony showing pyoverdine gene expression under stringent iron limitation (CAA + 400μM bipyridyl) used as a representative example. Branch length indicates the lifespan of a cell and color indicates the intensity of average pyoverdine gene expression of the respective cell during its lifespan. (**b**) Positive associations between the lifespans of cells and their siderophore gene expression (pyoverdine =blue, pyochelin=red) when not controlling for cell generation as a confounding factor. The panel shows data from the lineage tree in (a). The lines indicate the smoothed conditional means. (**c+d**) Partial correlation coefficients between the lifespan of cells and their siderophore gene expression, controlling for generation identity under moderate (CAA + 0μM bipyridyl) and stringent (CAA + 400μM bipyridyl) iron limitation. Each dot represents the partial correlation coefficient of a colony (excluding the cells on the tip of the trees, which did not divide during the duration of the assay). Boxplots show the median and the interquartile range (IQR), while whiskers indicate minimum and maximum values. Asterisks indicate significant differences from the dashed line. ** p < 0.01.

We indeed found that cell life span correlated positively with both pyoverdine and pyochelin gene expression (Fig. 3b shows a representative example). However, the lineage trees also revealed that cell division rates accelerated over time, such that the first generations of cells have longer life spans than the later generations of cells (Fig. 3a). To control for this confounding factor, we calculated the partial correlations between lifespans of cells and gene expression, whilst controlling for generation identity. We found no longer any positive associations, neither between pyoverdine gene expression and lifespan (moderate iron limitation, *t*_18_ = -1.85, p = 0.0807, stringent iron limitation, *t*_25_ = -0.65, p = 0.5200), nor between pyochelin gene expression and lifespan (under moderate iron limitation, the association is negative, *t*_18_ = -3.20, p = 0.0049, stringent iron limitation, *t*_25_ = 1.37, p = 0.1830). Thus, variation in cellular lifespan does not seem to contribute to siderophore gene expression heterogeneity.

### Sister cells have more similar siderophore gene expression patterns than neighbors

We hypothesized that epigenetic inheritance results in daughter cells showing more similar gene expression levels than non-related cells. When testing this hypothesis, it is important to consider that sister cells are often spatially next to one another after cell division on structured surfaces. Thus, sisters likely experience similar micro-environmental conditions, such that gene expression might be similar because they share the same environment and not because of shared genealogy. To disentangle environmental from genealogical effects, we used our lineage trees to identify sister cells and their closest neighbors originating from a different mother cell. On average, we analyzed 138 sister-sister vs. sister-neighbor pairs per colony and found that sister cells consistently showed more similar gene expression levels than unrelated neighbors (Fig. 4). Specifically, gene expression differences between sister cells were significantly smaller than gene expression differences between closest neighbors for both pyoverdine (Fig. 4a, F_1,86_ = 37.44, p< 0.0001) and pyochelin (Fig. 4b, F_1,86_ = 28.06, p < 0.0001) gene expression. The gene expression differences between sister cells and closest neighbors were similar between the two levels of iron limitation (pyoverdine: F_1,86_ = 0.023, p = 0.879; pyochelin: F_1,86_ = 0.134, p = 0.715). Hence, genealogy is an important determinant of siderophore gene expression heterogeneity.

**Fig 4.**
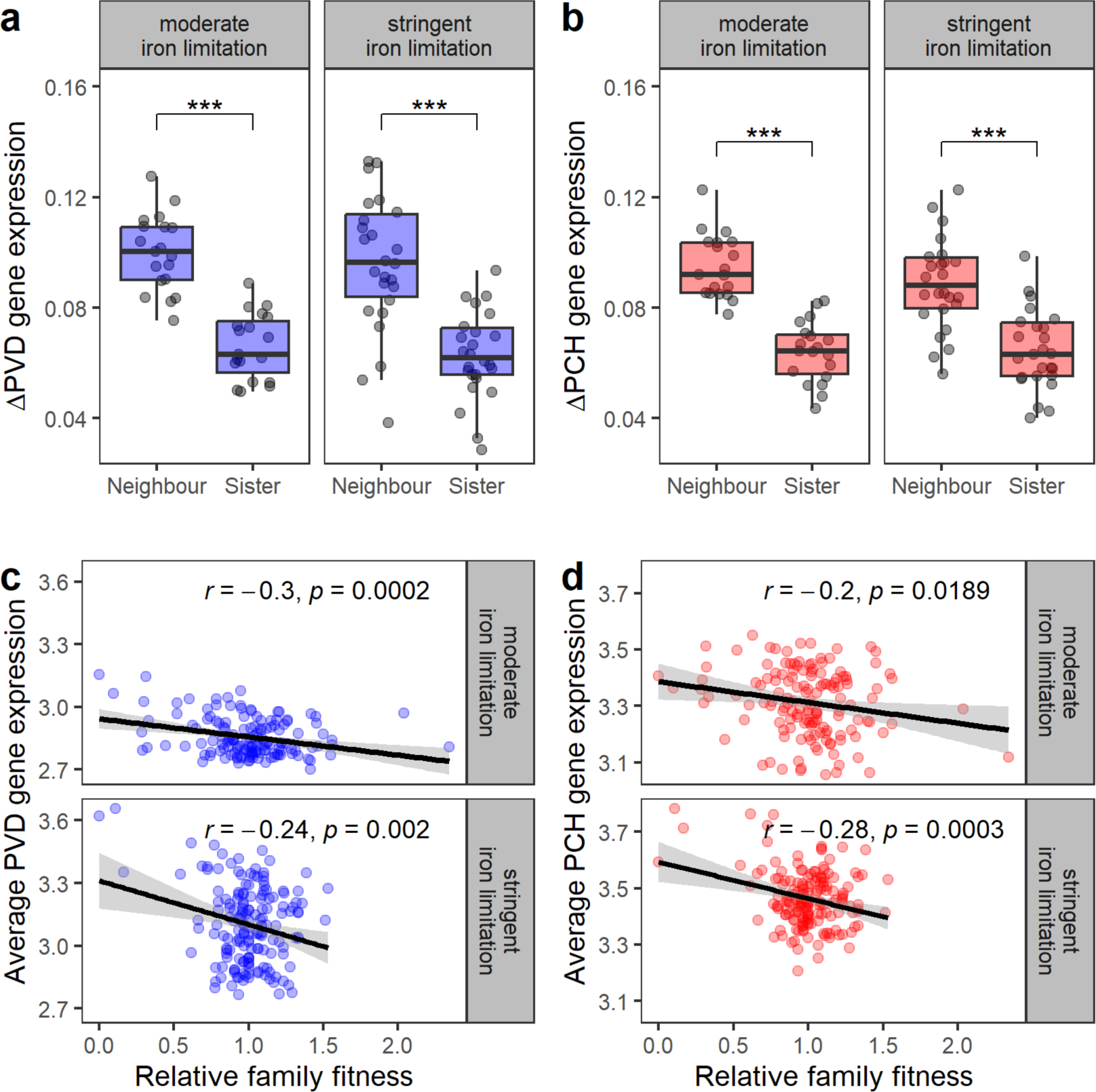
Sisters have more similar siderophore gene expression levels than closest neighbors. Comparison of gene expression differences between sister cells and closest neighbors for pyoverdine (**a**, blue, PVD) and pyochelin (**b**, red, PCH) gene expression. The difference in gene expression between all sister cell pairs and closest neighbor pairs (excluding sisters) was calculated. Data points represent mean differences per colony, for neighbors or sisters. Boxplots show the median and the interquartile range (IQR), while whiskers indicate minimum and maximum values, excluding outliers. Four outlier values (all from the same colony under stringent iron limitation, one each in the four categories in a) are not shown but were included in the statistical analyses. (**c**) Correlations between the number of offspring in a family branch and its average pyoverdine gene expression level. (**d**) Correlations between the number of offspring in a family branch and its average pyochelin gene expression level. For (c) and (d), each colony was split into four family branches and the fitness (number of cells) of each branch was calculated relative to the average within the colony, which was then contrasted against the average siderophore gene expression within the family branch. R-values indicate Pearson’s correlation coefficients. * p < 0.05, ** p < 0.01, *** p < 0.001.

Given that there are genealogical differences in siderophore gene expression among cell lineages within a colony, we asked whether siderophore over-expression or under-expression, relative to the colony average, is associated with fitness consequences. If siderophores primarily benefit the producer, one would expect cell lineages that over-produce siderophores to have more offspring. By contrast, if siderophores are evenly shared within the colony one would expect cell lineages that under-produce siderophores to have more offspring, because they save production costs whilst reaping equal benefits. To differentiate between the two scenarios, we split each colony into four family lineages (after the second cell division) and related the number of offspring in each family lineage to its average siderophore gene expression level. We found significant negative correlations between the two metrics under moderate and stringent iron limitations for pyoverdine (Fig. 4c) and pyochelin (Fig. 4d), lending support to the second scenario and thus a shared function of siderophores.

### Cell pole age does not correlate with siderophore gene expression

We hypothesized that variation in cell pole age could translate into gene expression heterogeneity across cells. To test this hypothesis, we used the DeLTA tracking algorithm ^31^ to infer cell pole age from the previously constructed lineage trees across three generations (Fig. S1). We then tested whether siderophore gene expression is associated with cell pole age. We found no support for our hypothesis, as there was no association between cell pole age and siderophore gene expression, neither for pyoverdine (Fig. 5a; F_5,258_ = 0.186, p = 0.9676) nor for pyochelin (Fig. 5b; F_5,258_ = 0.828, p = 0.5304).

**Fig 5.**
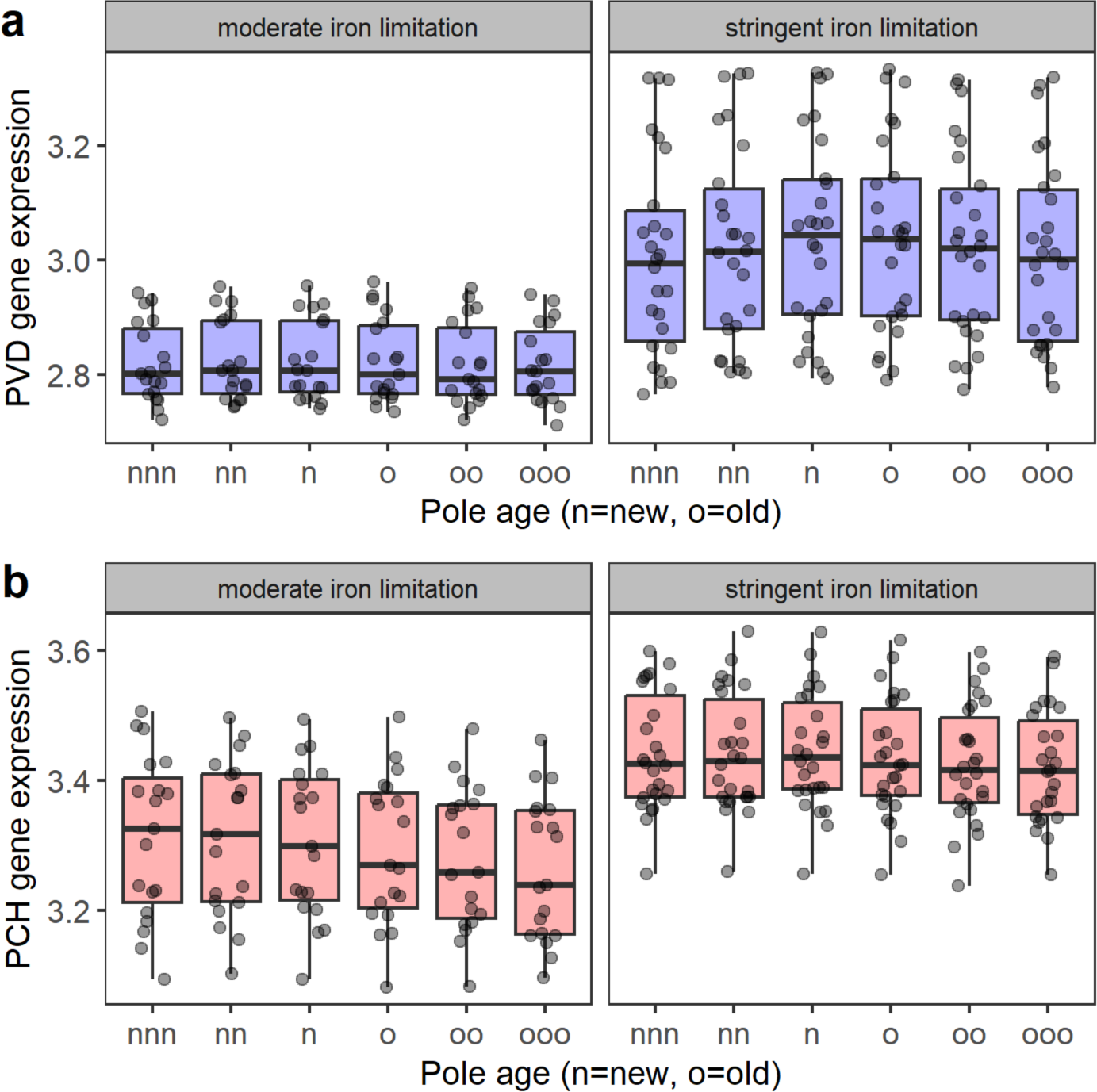
Cell pole age does not correlate with siderophore gene expression. (**a**) Pyoverdine and (**b**) pyochelin gene expression in response to cell pole age under moderate (CAA + 0μM bipyridyl) and stringent iron limitation (CAA + 400μM bipyridyl). Cells are categorized based on the cell pole they inherited from their mother (n or o, first letter), as well as the pole their mothers (nn or oo, second letter) and grandmothers (nnn or ooo, third letter) received. The analysis only considered ‘pure’ cousins (nn and oo) and second cousins (nnn and ooo), whereas cells with mixed cell poll age histories (e.g., nno) were not included. Individual dots show the mean gene expression of the corresponding pole age category per colony. Boxplots show the median and the interquartile range (IQR), while whiskers indicate minimum and maximum values.

## Discussion

We set out to identify extrinsic and intrinsic sources of cellular heterogeneity in the expression of pyoverdine and pyochelin synthesis genes in the bacterium *Pseudomonas aeruginosa* and relate them to adaptive (evolutionary) functions. The siderophores pyoverdine and pyochelin are secreted in the environment and shared as public good between cells^30^, meaning that heterogeneity in siderophore investment can have fitness consequences for the producing individual and the other group members. When grown on surfaces, we found that the spatial positioning of cells within the colony and cell lineage genealogy were major sources of cellular heterogeneity. By contrast, cellular lifespan and cell pole age had no significant effect on heterogeneity. At the functional level, our results suggest that bacteria optimize iron acquisition strategies depending on their location in the colony. Optimization was particularly apparent under moderate iron limitation, under which edge cells predominantly invested in the potent yet expensive pyoverdine, while interior cells specialized on the production of the cheaper, less potent pyochelin. We further observed negative correlations between siderophore gene expression of cell lineages within colonies and their fitness, suggesting that siderophore-overproducing cell lineages boost the fitness of siderophore-underproducing lineages through siderophore sharing. Altogether, our study shows that a combination of single-cell time-lapse microscopy together with quantitative image analysis is a powerful tool to reveal mechanistic and adaptive causes of cellular heterogeneity in clonal bacterial populations.

We observed spatial gradients of siderophore gene expression activities from the center to the edge of colonies (Fig. 2). Gradients were more prominent in larger colonies and predominantly occurred under moderate iron limitation. At the mechanistic level, these results indicate that bacteria can sense small differences in iron availabilities. Indeed, *P. aeruginosa* can adjust siderophore production in a remarkably fine-tuned manner. Upon the depletion of intra-cellular iron stocks, Fur (ferric uptake regulator) loses its inhibitory effect, resulting in the basal expression of siderophore synthesis enzymes^21,32^. Pyoverdine and pyochelin are both produced via non-ribosomal peptide synthesis and are actively secreted via specific exporters^21,33^. Production levels are then fine-tuned based on incoming iron-loaded siderophores triggering positive feedback loops. For pyoverdine, the signaling cascade involves an interplay between the sigma factor PvdS and its antagonist FpvR^23,34^. For pyochelin, signaling is based on a direct interaction between ferri-pyochelin and the transcriptional regulator PchR^24,35^. A fine-tuned response is achieved by the relative strength of FUR-mediated repression and signaling-mediated activation. While pyoverdine and pyochelin are co-regulated overall^19^, preferential pyochelin production can occur under moderate iron limitation because FUR repression seems to be more relaxed for this siderophore^28^. Conversely, preferential pyoverdine production can occur under stringent iron limitation, most likely because pyoverdine inhibits pyochelin-mediated signaling as it has stronger affinity for iron^19,27,28^. At the evolutionary level, our results suggest that bacteria optimize their iron acquisition strategy under moderate iron limitation, depending on their position in the colony as captured by our adaptive explanation A. In larger colonies, center cells might predominantly recycle the pyoverdine that has already been produced by earlier generations and thus switch to the production of the cheaper pyochelin. By contrast, edge cell might still experience a shortage of siderophores and iron, and therefore predominantly invest in the more potent pyoverdine. Compatible with previous work, this optimization only occurs under moderate iron limitation, conditions under which pyochelin becomes a potent siderophore^28^.

Besides spatial effects, we found that epigenetic inheritance is a major factor explaining cellular heterogeneity in siderophore gene expression. Daughter cells inherit the gene expression status from their mother and differences between families are propagated through the lineage tree. Similar observations were made for the expression of several genes in *Escherichia coli*^9^. While patterns of epigenetic inheritance are consistent across genes and levels of iron limitation (Fig. 4), the genealogical effects do not explain how variation in gene expression across cell lineages arises in the first place. One option is that random noise in gene expression leads to variation among the first few founder cells in a colony and this variation is then passed on to all the subsequent lineage members. However, the fact that random noise is expected to dilute rather than propagate genealogical differences speaks against this hypothesis. Alternatively, we have previously shown that siderophore gene expression heterogeneity is positively linked to the metabolic activity of cells^19^. Accordingly, cells could therefore differ in their vigour^7^, whereby cells with relatively low vigor invest less in siderophores than cells with relatively high vigor, and the vigor status is inherited from mother to daughter cells. Based on our data, we consider this as a likely explanation that would need to be experimentally substantiated in the future.

Important to note is that pyoverdine and pyochelin are diffusible molecules that serve as signals when taken up by cells in their iron-bound forms^23,24^. This means that siderophores produced by one cell can induce siderophore production in a neighboring cell. In a spatial setting, siderophore diffusion and signaling are expected to affect cellular heterogeneity. For example, molecule diffusion is limited in structured environments^17,36,37^ such that siderophore-mediated signaling should primarily occur between neighboring cells. Local signaling should strengthen spatial correlations. This is indeed supported by our data showing that spatial correlations predominantly arise in larger colonies (Fig. 2b), in which signaling is expected to be more intense and local. These spatial effects can also be seen at a qualitative (visual) level in Fig. 2a, where we observe patches of cells with above- and below-average gene expression, patterns that can be induced by high and low local signaling, respectively. In contrast, local siderophore signaling is expected to weaken the genealogical effects (Fig. 4a+b) since diffusing siderophores should induce siderophore production in both sister and unrelated neighbors.

We found no support for the adaptive explanation B, namely that cells with low individual fitness can indirectly increase group fitness via disproportionally high siderophore investment levels. There was no support for this hypothesis when relating cell life span to siderophore gene expression (Fig. 3). Cells differ in their lifespan, but slower dividing cells did not have higher siderophore investment than faster dividing cells. There was also no support for our hypothesis when focusing on cell pole age, as this metric showed no association with siderophore gene expression levels (Fig. 5). However, we unexpectedly observed a link between epigenetic inheritance and fitness, whereby cell lineages with below-average siderophore investment levels had above-average cell division rates (Fig. 4). Building on our vigor model, this result indicates that cell lineages with high vigor produce high amounts of siderophore to boost not only their own fitness but also the fitness of low-vigor cell lineages trough the cooperative sharing of siderophores.

In conclusion, we conducted simple time-lapse microscopy experiments to track gene expression of individual bacterial cells and cell lineages in growing colonies on surfaces over time. Together with automated image analysis and cell-tracking software, we show that an enormous amount of information can be extracted from such simple experiments. The approach chosen in our paper does not only allow to test hypotheses regarding the sources of cellular gene expression heterogeneity but also to examine potential fitness consequences. Taken together, we advocate that future bacterial single-cell studies should have a strong focus on fitness aspects to examine to what extent gene expression heterogeneity reflects random noise as opposed to exerting an adaptive function.

## Materials and methods

### Bacterial strains

For all our experiments, we used the standard laboratory strain *P. aeruginosa* PAO1, which produces the siderophores pyochelin and pyoverdine. To measure gene expression, we used PAO1 strains with fluorescent gene reporter constructs chromosomally integrated as single copies, at the *att*Tn7 site of the wild type using the mini-Tn7 system^38^ and our customized protocols^39^. To simultaneously track the expression of two genes, we used a double gene expression reporter: PAO1*pvdA*::*mcherry*–*pchEF*::*gfp*, in which the promoter for the pyoverdine biosynthetic gene *pvdA* is fused to the red fluorescent gene *mcherry* and the pyochelin biosynthetic genes *pchE* and *pchF* (forming an operon) are fused to the green fluorescent gene *gfpmut3*. For simplicity of nomenclature the *gfpmut3* is referred to as *gfp*. The genetic scaffold of the double gene reporter construct, and its construction is described in detail elsewhere^19^. In this earlier study, we have also demonstrated that gene expression levels correlate well with the actual amount of siderophores produced.

### Growth conditions

Prior to experiments, we prepared overnight cultures from -80 °C stocks, in 8ml Lysogeny broth (LB) in 50ml tubes, incubated at 37°C, shaken at 220 rpm for approximately 18 hours. Cells were then harvested by centrifugation (5000 rpm for 3 minutes), subsequently washed in 0.8% saline, and adjusted to OD_600_ = 0.001 (optical density at 600nm). To stimulate a substrate attached mode of growth, harvested cells were then seeded onto 1% agarose pads on a microscopy slide. The medium of the pad consisted of CAA (5g casamino acids, 1.18g K_2H_PO_4*_3H_2O_, 0.25g MgSO_4*_7H_2O_, per litre), buffered at physiological pH by the addition of 25mM HEPES. To induce a gradient of iron limitation, we used either plain CAA or CAA supplemented with 400µM of the synthetic iron chelator 2-2’-bipyridyl. All chemicals were purchased from Sigma Aldrich (Buchs SG, Switzerland). The gene expression of individuals within growing colonies was quantified using a widefield fluorescence microscope (Olympus ScanR), featuring an incubation chamber where cells were incubated at 37 °C for 5 hours.

### Preparation of microscope slides

To prepare agarose pads on microscopy slides, we adapted a method previously described elsewhere^19,40,41^. Standard microscope slides (76mm x 26mm) were sterilized with 70% ethanol. We used ‘Gene Frames’ (Thermo Fisher Scientific, Vernier, Switzerland) to prepare agarose pads on which bacteria were seeded. Each frame features a single chamber (17 mm x 28 mm) of 25 mm thickness. The frames are coated with adhesives on both sides so that they stick to the microscope slide and the coverslip. The sealed chamber is airtight, which prevents pad deformation and evaporation during experimentation.

To prepare agarose pads, we heated 40 ml of plain CAA medium with 1% agarose in a microwave. The agarose-CAA solution was first cooled to approximately 50°C. Then 25mM HEPES buffer along with the required concentration of bipyridyl was added. We pipetted 700 µl of the solution into the gene frame and immediately covered it with a sterile coverslip. The coverslip was gently pressed to let superfluous medium escape and solidify for around 20 minutes. After solidification, we removed the coverslip (by carefully sliding it sideways) and divided the agarose pad into four smaller pads of roughly equal size with a sterile scalpel. We introduced channels between pads, that served as oxygen reservoirs for the growing colonies. To ensure that colonies started to grow from a single cell, we put 1 µl of diluted bacteria (OD_600_ = 0.001) on each agarose pad. Upon the addition of bacteria, we let the agarose pads air-dry for 2 minutes, and then sealed them with a new sterile coverslip.

### Microscope set-up and time-lapse imaging

Following the preparation steps described above, we immediately started time-lapse imaging of the bacteria at the Center for Microscopy and Image Analysis of the University of Zurich (ZMB) using an inverted widefield Olympus ScanR HCS microscope, featuring an incubation chamber. The microscope has an automatic movable stage, capable of imaging multiple fields of view repeatedly and a motorized Z-drive, which enables autofocusing of the objects. The microscope is controlled by the OLYMPUS cellSens Dimensions software. Images were captured with a PLAPON 60x phase oil immersion objective (NA=1.42, WD=0.15mm) and a Hamamatsu ORCA_FLASH 4.0V2, high sensitive digital monochrome scientific cooled sCMOS camera (resolution: 2048x2048 pixels, 16-bit).

For time-lapse microscopy, we first imaged the growing colonies with phase contrast (exposure time 56.4 ms). For fluorescence imaging, we used a fast emission filter wheel, featuring a FITC SEM filter for GFP (exposure time 50 ms, excitation=470±24 nm, emission=515±30nm, DM=485) and a TRITC SEM filter for mCherry (exposure time 50 ms, excitation=550±15nm, emission=595±40nm, DM=558). Imaging with phase contrast and the respective fluorescent channels was done sequentially for every time point and field of view. The time-lapse image recording was performed at 37 °C for 5 hours with images taken every 10 minutes. We started the time-lapse image recording with a field of view having a maximum of three separate cells.

### Image processing, single cell segmentation and tracking

We used FIJI^42^, Ilastik^43^ and DeLTA^31^ to (i) process images, (ii) segment single cells, (iii) track lineages and (iv) measure fluorescence in our time-lapse recordings. We conducted a preliminary quality check by inspecting all time-lapses in FIJI. We removed recordings that were blurry, had excessive drift, and cases in which cells began to grow in double layers. Recordings of two positions without cells were used to create average blank images for the two fluorescence channels. These average blank images were then subtracted from all fluorescence images to correct for microscope vignetting across the fields of view. Thereafter, we performed a drift correction of our time-lapse recordings (https://github.com/fiji/Correct_3D_Drift) and exported the images for segmentation into Ilastik.

We segmented the single cells based on the phase contrast images with the pixel and object classification workflow in Ilastik version 1.3.2^43^: The classifier was trained with a random sample of 12 images from our collection. Subsequently, the bulk of images were segmented in batch mode. Based on the Ilastik segmentations, we create regions of interest (ROI) for individual cells using FIJI. To correct for background fluorescence, we measured the mean fluorescence intensity outside the ROIs for each image and subtracted the corresponding value from the image.

We used DeLTA, a segmentation and tracking pipeline based on deep convolutional neural networks. We used this platform for cell identity and lineage tracking for all our time-lapse recordings. The workflow also involves segmentation, which allowed us to compare segmentation results from DeLTA and Ilastik. We found that segmentation with DeLTA leads to slightly smaller cells, compared to Ilastik segmentation (Fig. S3a+b). To validate automated cell tracking, we manually tracked cell lineages for eight colonies using the segmentation from DeLTA. We found that tracking accuracy decreased at later time points, but cell and division tracking accuracy were still ≥ 95% for seven out of eight and five out of eight colonies, respectively (Tab. S3).

### Single-cell measurements and data analysis

We extracted information on each cell’s position, size, and fluorescence intensity in the two fluorescent channels (GFP and mCherry) with FIJI. The subsequent analyses were conducted using R (4.3.0). The single cell data allowed us to determine the number of cells present at each timepoint of a recording. For positions where the time-lapse recording started with one single cell, the number of cells at each timepoint gave us the colony size over time. In positions that started with multiple single cells scattered over the field of view, we grouped cells into individual colonies with a hierarchical cluster analysis of the distance between cells. To quantify siderophore gene expression, we generally used the integrated density of the fluorescence intensity. There was one exception: we used the mean fluorescence intensity when analyzing expression correlations between the two siderophore genes. This is because the integrated density correlates with cell size, such that positive correlations necessarily arise when there is variation in cell size. We applied a log_10 t_ransformation to both types of fluorescence intensity values.

For colony-level analysis, we calculated the average siderophore gene expression intensity across cells within a colony per timepoint. These values were then regressed over time to test whether average siderophore gene expression changes over time (positive or negative slope). We further analyzed correlations between the expression of the two siderophore genes, including all colonies with at least 16 cells or larger, and assessed whether the correlations change over time.

For single-cell analyses, we determined the position of each cell within a colony by the distance of a cell to the edge of the colony. For each time-point, we identified the edge of the colony by computing the α-convex hull for the colony with an alpha value of 30 pixels, which corresponds approximately to the length of a cell. This approach allowed us to identify the cells that were on the edge of a colony. Connecting the centers of these cells gave us the outer contour of a colony. Subsequently, we calculated the Euclidean distance between each cell’s centre and its distance to the nearest edge of the colony. For each colony, we then calculated the spatial correlation between siderophore gene expression and distance to the edge across cells per timepoint. For statistical comparisons, we grouped the colonies into four size categories ([16,31], [32,63], [51,127], [128,256]), excluding colonies with less than 16 or more than 256 cells.

We used the results from the DeLTA pipeline for three different analyses involving cell lineage tracking. First, we calculated the lifespan of each cell, defined as the number of timepoints between two consecutive cell divisions. This metric was then correlated with the average siderophore gene expression intensity over the lifespan of cells. Given that lifespans decreased in later generations, we conducted partial correlation analyses, controlling for the generation number a cell belongs to. Second, we identified a sister and a closest neighbor cell for each cell in a colony. Sister cell pairs could be directly derived from the lineage tree. To identify closest neighbors, we first calculated the average position (coordinates) of cells over their lifespans. We then determined closest neighbors as pairs of cells with the shortest distance between their average positions that were born at most one timepoint apart, and that were not sisters. This analysis was conducted for all colonies with four and more cells. To avoid double counting, the analysis was restricted to only one cell within sister pairs. Third, we tracked the cell pole age for each cell across three generations. Upon cell division, each daughter cell inherits an old pole from its mother and forms a new pole (Fig. S1). DeLTA records pole age so that cells can be classified as “n” (new) or “o” (old) pole cells considering the past cell division, and as “nn” or “oo” and “nnn” or “ooo” cells when further considering division events of their mothers and grandmothers, respectively. For simplicity reasons, we did not include cells with mixed cell poll age histories.

### Statistical analysis

We used general linear models for statistical analysis in R 4.3.0. Prior to analysis, we used the Shapiro-Wilk test to confirm that model residuals are normally distributed. For colony-level analysis (Fig. 1), we used analysis of variance (ANOVA) models to assess whether pyoverdine and pyochelin gene expression and the correlation between the two vary in response to iron limitation. We further used two-sided one-sample *t*-tests to test for significant temporal changes of these three variables (i.e. correlations coefficients being different from zero). At the single-cell level, we used two-way ANOVAs to analyze whether spatial correlation of siderophore gene expression differ in response to colony size and iron limitation (Fig. 2). Subsequently we used two-sided one-sample *t*-tests to determine whether spatial correlation coefficients across colonies are significantly different from zero. Reported P-values were adjusted for multiple testing using the Holm-Bonferroni method. To analyze whether cell lifespan relates to pyoverdine and pyochelin gene expression, we calculated the partial correlation coefficients across cells separately for each colony, whilst controlling for the generation number each cell belongs to (Fig. 3). Subsequently, we used two-sided one-sample *t*-tests to test whether the partial correlation coefficients differ from zero. We used two-way ANOVAs to test whether siderophore gene expression is more similar between sister cells than between unrelated neighbors and whether differences depend on the level of iron limitation. To test if there is an association between the average siderophore gene expression and the number of offspring in a family branch we calculated the Pearson’s correlation coefficient. To analyse whether cell pole age influences siderophore gene expression, we used two-way ANOVAs with pole age category and iron limitation as fixed factors (Fig. 5).

## Acknowledgments

We thank Priyanikha Jayakumar for her help in the lab, and the Center for Microscopy and Image Analysis for support. This project has received funding from the European Research Council (ERC) under the European Union’s Horizon 2020 research and innovation programme (grant agreement no. 681295).

## Author contributions

SM and RK designed the study. SM carried out all experiments. TW wrote the image analysis scripts. SM, TW and RK analyzed the data and wrote the paper.

