## Supplementary Figures and Tables for "Space and epigenetic inheritance determine inter-individual differences in siderophore gene expression in bacterial colonies"

This file contains the following supplementary materials:

- 3 supplementary tables
- 3 supplementary figures

**Tab S1.** Difference of special correlation PVD to zero from Fig 2b with corresponding p-values

| Size category | Media | df | t | p |
|---|---|---|---|---|
| [16,31] | 0µM bipyridyl | 38 | -4.2988 | 0.0007 |
| [32,63] | 0µM bipyridyl | 69 | -7.1432 | < 0.0001 |
| [64,127] | 0µM bipyridyl | 57 | -6.8388 | < 0.0001 |
| [128,256] | 0µM bipyridyl | 44 | -3.7052 | 0.0029 |
| [16,31] | 400µM bipyridyl | 91 | -1.2835 | 0.2030 |
| [32,63] | 400µM bipyridyl | 77 | -1.6585 | 0.2020 |
| [64,127] | 400µM bipyridyl | 55 | 2.6349 | 0.0436 |
| [128,256] | 400µM bipyridyl | 16 | 2.6719 | 0.0501 |

**Tab S2.** Difference of special correlation PCH to zero from Fig 2b with corresponding p-values

| Size category | Media | df | t | p |
|---|---|---|---|---|
| [16,31] | 0µM bipyridyl | 38 | -0.8246 | 1.0000 |
| [32,63] | 0µM bipyridyl | 69 | 2.0869 | 0.1624 |
| [64,127] | 0µM bipyridyl | 57 | 4.0015 | 0.0011 |
| [128,256] | 0µM bipyridyl | 44 | 6.1068 | < 0.0001 |
| [16,31] | 400µM bipyridyl | 91 | -0.3280 | 1.0000 |
| [32,63] | 400µM bipyridyl | 77 | -0.1487 | 1.0000 |
| [64,127] | 400µM bipyridyl | 55 | 4.5324 | 0.00022 |
| [128,256] | 400µM bipyridyl | 16 | 3.3498 | 0.02035 |

**Tab S3.** DeLTA tracking and division accuracy in percent up to the 10th frame, the 20th frame, and the last frame for 8 manually tracked positions.

| Position | Final number of cells | Number of frames | Correct tracking frame 10 | Correct tracking frame 20 | Correct tracking last frame | Correct divisions frame 10 | Correct divisions frame 20 | Correct divisions last frame |
|---|---|---|---|---|---|---|---|---|
| Bp000 _1 | 188 | 30 | 1 | 0.98 | 0.99 | 1 | 0.98 | 0.97 |
| Bp000 _2 | 129 | 30 | 1 | 1 | 0.99 | 1 | 1 | 0.98 |
| Bp000 _4 | 571 | 29 | 1 | 1 | 0.86 | 1 | 1 | 0.72 |
| Bp000 _8 | 360 | 30 | 1 | 0.99 | 0.98 | 1 | 0.97 | 0.94 |
| Bp400 _2 | 267 | 30 | 1 | 1 | 0.98 | 1 | 1 | 0.95 |
| Bp400 _3 | 282 | 30 | 1 | 1 | 0.98 | 1 | 1 | 0.95 |
| Bp400 _1 | 240 | 27 | 1 | 1 | 0.95 | 1 | 1 | 0.90 |
| Bp400 _3 | 142 | 29 | 1 | 1 | 0.99 | 1 | 1 | 0.97 |

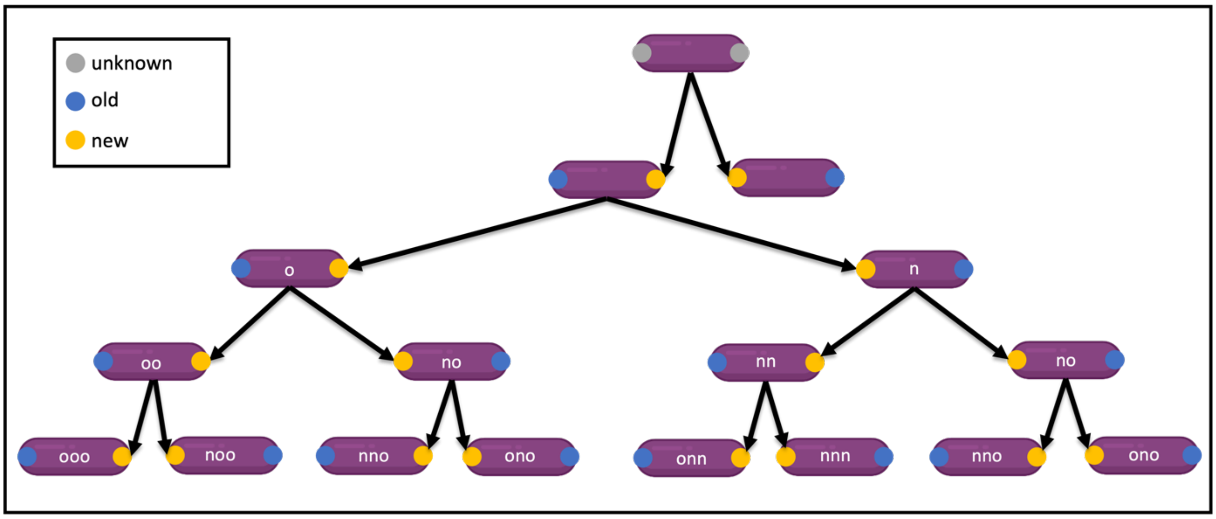


**Fig S1.** Pole inheritance scheme. The DeLTA pipeline determines which of the two poles of a cell is new, created by the division of the mother cell, or old. A cell is classified as having a new pole (n) if it receives the new cell pole from its mother. For the classification of subsequent generations, the new classification is appended to the front. The category “no” for example is assigned to a cell that received the new pole from its mother. The mother previously received an old pole from the grandmother.

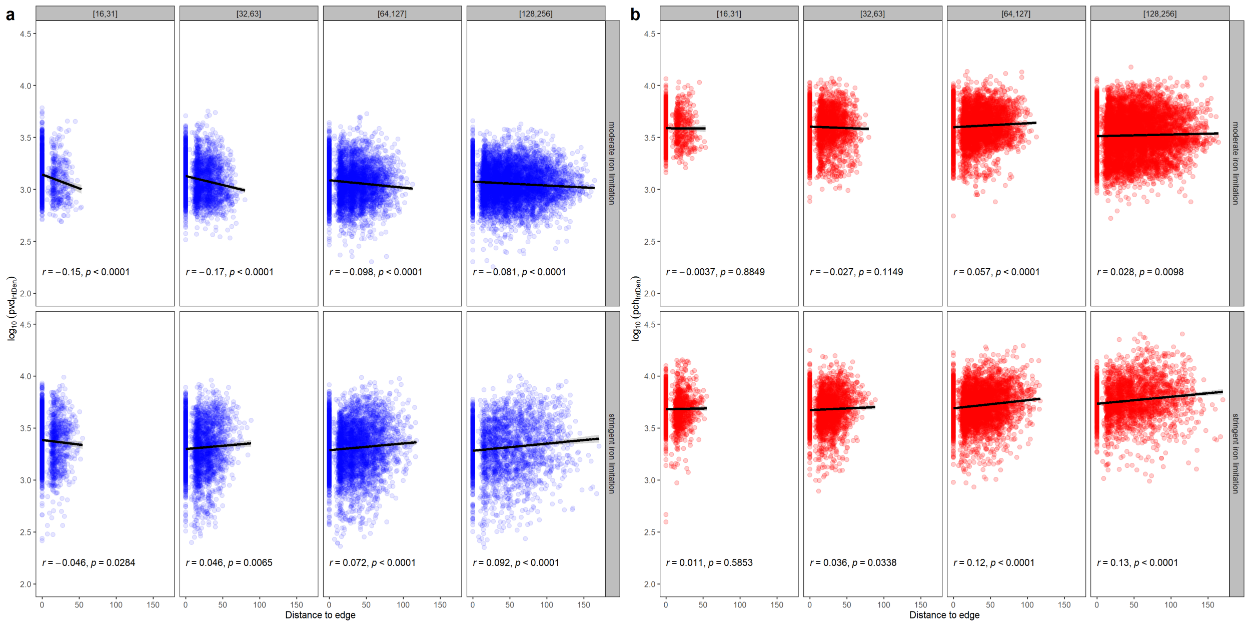

**Fig S2.** Siderophore gene expression of (a) pyoverdine (b) pyochelin over distance to the edge of the colony, for the different colony size categories. Black lines indicate the linear fit to the data, *r* the Pearson correlation coefficient together with the *p*-value of the fit.

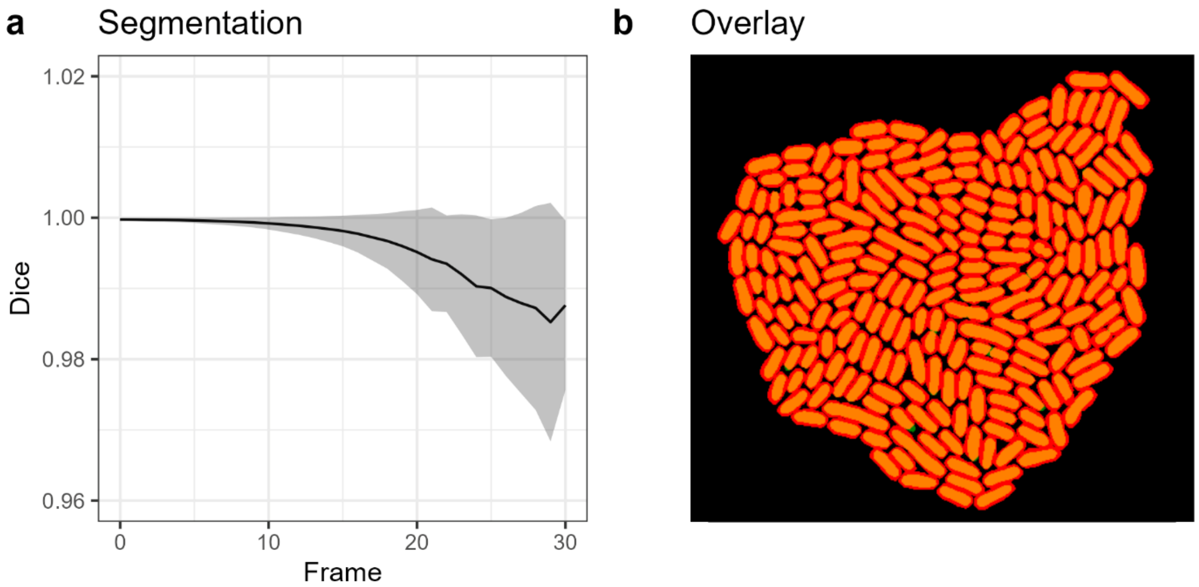


**Fig S3** Metrics to assess the accuracy of the segmentation from the DeLTA pipeline. **(a)** The dice score between the Ilastik segmentation and the DeLTA segmentation. The Dice score is calculated based on the formula DSc = 2TP/(2TP+FP+FN), where TP is the number of overlapping pixels, FP is the number of pixels wrongly segmented as cells, and FN is the number of pixels wrongly segmented as background (FN). **(b)** Representative example of the overlap between the Ilastik segmentation and the DeLTA segmentation. Orange is where the two segmentations overlap, red where Ilastik allocated pixels to a cell but not DeLTA, and green where DeLTA allocated pixels to a cell but not Ilastik. The comparison shows that there is high overlap and consistency between the two platforms, but cells are overall larger with Ilastik.
